# Insights into the mutation T1117I in the spike and the lineage B.1.1.389 of SARS-CoV-2 circulating in Costa Rica

**DOI:** 10.1101/2021.07.08.451640

**Authors:** Jose Arturo Molina-Mora

## Abstract

Emerging mutations and genotypes of the SARS-CoV-2 virus, responsible for the COVID-19 pandemic, have been reported globally. In Costa Rica during the year 2020, a predominant genotype carrying the mutation T1117I in the spike (S:T1117I) was previously identified. To investigate the possible effects of this mutation on the function of the spike, i.e. the biology of the virus, different bioinformatic pipelines based on phylogeny, natural selection and co-evolutionary models, molecular docking and epitopes prediction were implemented.

Results of the phylogeny of sequences carrying the S:T1117I worldwide showed a polyphyletic group, with the emergency of local lineages. In Costa Rica, the mutation is found in the lineage B.1.1.389 and it is suggested to be a product of positive/adaptive selection. Different changes in the function of the spike protein and more stable interaction with a ligand (nelfinavir drug) were found. Only one epitope out 742 in the spike was affected by the mutation, with some different properties, but suggesting scarce changes in the immune response and no influence on the vaccine effectiveness.

Jointly, these results suggest a partial benefit of the mutation for the spread of the virus with this genotype during the year 2020 in Costa Rica, although possibly not strong enough with the introduction of new lineages during early 2021 which became predominant later. In addition, the bioinformatics pipeline offers an integrative and exhaustive *in silico* strategy to eventually study other mutations of interest for the SARS-CoV-2 virus and other pathogens.

**Highlights:**

- In Costa Rica during the year 2020, a predominant SARS-CoV-2 genotype carrying the mutation T1117I in the spike (S:T1117I) was identified.
- The S:T1117I was assessed for possible effects of this mutation on the function of the spike with a *in silico* approach.
- Phylogeny revealed that sequences carrying the S:T1117I worldwide define a polyphyletic group, with the emergency of local lineages, including the lineage B.1.1.389 in Costa Rica.
- A positive/adaptive selection was identified for S:T1117I, with different changes in the function of the spike protein, more stable interaction with ligands and scarce changes in the immune response.
- The bioinformatics pipeline can be eventually used to study other mutations of the SARS-CoV-2 virus and other pathogens.

## Introduction

The SARS-CoV-2 virus, responsible for the COVID-19 pandemic, has been detected in more than 181 million people worldwide and more than 365 000 cases in Costa Rica (with at least 4650 deaths) until June 30^th^, 2021. As part of the epidemiological surveillance of the pandemic in Costa Rica, we have studied both genomic features of the virus and clinical and demographic patterns among diagnosed patients (Molina-Mora, Cordero-Laurent, et al., 2021; Molina-Mora, González, et al., 2021).

Emerging mutations in the spike and genotypes have been reported globally, including variants of interest (VOI) and variants of concern (VOC), which are still under study owed to possible or confirmed changes in transmission, severity, clinical manifestations, mortality, or vaccine effectiveness (Graham et al., 2021). In Costa Rica, during 2020 the predominant SARS-CoV-2 genome was a genotype carrying the mutation T1117I in the spike (S:T1117I), as we reported previously (Molina-Mora, Cordero-Laurent, et al., 2021). This local genotype, now classified as a Costa Rican PANGOLIN lineage B.1.1.389, reached up to 30% of cases in this country by December 2020 (Molina-Mora, Cordero-Laurent, et al., 2021). Furthermore, this lineage was distributed among all the distinct clusters or clinical profiles from Costa Rican cases of COVID-19, as we analyzed using machine learning (Molina-Mora, González, et al., 2021). For other geographic locations, other lineages carrying the mutation S:T1117I have been reported but, unlike Costa Rica, with a frequency <1% (https://www.gisaid.org/).

The growing interest in the spread of SARS-CoV-2 genotypes, including the VOC/VOI worldwide and the lineage B.1.1.389 in Costa Rica, is mainly explained by the mutations in the spike protein. Spike protein is relevant not only because of its interaction with the receptor in human cells (ACE2, angiotensin-converting enzyme 2) (Zhou et al., 2020), but also it is implicated in the activation of the immune response against the virus by natural infection or vaccination (Graham et al., 2021; Koyama, Platt, & Parida, 2020). Hence, changes in the spike protein sequence could affect the transmission and clinical manifestations (Graham et al., 2021; Toyoshima, Nemoto, Matsumoto, Nakamura, & Kiyotani, 2020), as well as the vaccine effectiveness (Koyama et al., 2020), i.e., the biology of the virus.

The spike protein is composed of 1273 amino acids and has two domains S1 (aa14–685) and S2 (aa686–1273) that are responsible for the binding step to the ACE2 receptor. The recognition and binding to receptors are part of the activity of the S1 domain, specifically with the RBD (receptor binding domain, aa319–541). A subsequent conformational change of the S2 domain, which harbors the putative fusion PF peptide (aa788–806) and the heptad repeats HR1 (aa912–984) and HR2 (aa1163–1213), facilitates the fusion between the envelope protein of the SARS-CoV-2 and the plasma membrane of the host cell (Astuti & Ysrafil, 2020; Xia et al., 2020).

In the case of the mutation S:T1117I, despite this is located between the HR1 and HR2 regions, predictions suggested possible effects on viral oligomerization needed for cell infection, and more studies were demanded to investigate the possible changes in transmissibility, severity, or vaccine effectiveness (Molina-Mora, Cordero-Laurent, et al., 2021). To this end, this work aimed to study in-depth the biological effects of the mutation S:T1117I on the function of the spike protein. To address that, different bioinformatic pipelines and mathematical models were implemented to assess the phylogenetic relationships among genotypes, possible natural selection in the local spread, co-evolutionary models, interaction with molecules, as well as an immune activity using genome sequencing data. In addition, the pipeline offers an integrative and exhaustive *in silico* strategy to eventually study mutations of interest for SARS-CoV-2 virus and other pathogens.

## Methods

### SARS-CoV-2 sequences

All the available SARS-CoV-2 sequences with the mutation S:T1117I, up to April 30^th^ and from all the locations worldwide, were retrieved from the GISAID database (Global Initiative on Sharing All Influenza Data, www.gisaid.org). Details for each genome is summarized in the Supplementary Material. The same database was used to obtain all the sequences from Costa Rica during the same period, in which the genomic region of the spike gene (ORF coordinates of the NC_045512.2 reference genome) was selected for each sequence using *getfasta*-Bedtools package (Quinlan & Hall, 2010). Epidemiological and genomic data for all the sequences by country or lineage were summarized using the Outbreak.info tool (https://outbreak.info/).

### Phylogenetic analysis of sequences with the mutation S:T1117I

All the worldwide sequences with the mutation S:T1117I were aligned using MAFFT v7.471 (Katoh, Misawa, Kuma, & Miyata, 2002). The construction of the phylogenetic tree model was achieved with IQ-TREE v1.6.12 (Minh et al., 2020), including ModelFinder (Kalyaanamoorthy, Minh, Wong, Von Haeseler, & Jermiin, 2017) to select the best nucleotide substitution model (using the Bayesian Information Criterion BIC, the best model was TN+F+I). Visualization was done using the iTOL tool v4 (Letunic & Bork, 2019).

### Positive selection analysis of mutations

To detect sites in the spike protein of positive/adaptive (as well as negative/purifying) selection, we performed an analysis of natural selection using HyPhy (hypothesis testing using phylogenies) (Kosakovsky Pond, Frost, & Muse, 2005). To this end, the Bayesian inference FUBAR (Fast, Unconstrained Bayesian AppRoximation) model (Murrell et al., 2013) was implemented using the alignment of spike sequences. Thus, we calculated the ratio of non-synonymous to synonymous substitutions (dN/dS) values, under the assumption that each variant appeared *de novo* in each individual, to define if the substitution was a product of a positive (dN/dS>1) or negative (dN/dS <1) selection. More details in (Ferrareze et al., 2021; Lythgoe et al., 2021).

### Coevolutionary analysis of spike sequences

All the spike sequences from SARS-CoV-2 genomes from Costa Rica were aligned using MAFFT v7.471 (Katoh et al., 2002). The reference sequence was also included. Subsequently, the MISTIC2 tool with default parameters (Colell, Iserte, Simonetti, & Marino-Buslje, 2018; Simonetti, Teppa, Chernomoretz, Nielsen, & Marino Buslje, 2013) was used to assess co-evolutionary couplings and to identify functionally important residues in the spike protein, using mutual information and residues conservation models. The structural model of the spike (see Molecular Docking section) was incorporated to predict effects on the structure.

### Molecular docking

To assess the effects of the mutations S:D614G and S:T1117I, the two spike mutations of the lineage B.1.1.389, on the interaction of the spike with a specific ligand, we performed a molecular docking analysis. For the ligand selection, because the 1117 position is between the HR1 and HR2 regions of the spike, we selected the nelfinavir drug which is known to bind spike in the HR1 region (Musarrat et al., 2020).

The structural model of the spike (ID: 6VXX) was obtained from PDB (https://www.rcsb.org/), and the molecular structure of the nelfinavir drug, which is known, was downloaded from Drugbank (https://go.drugbank.com/). The analysis was done using the reference spike PDB model (wild type, WT) as well as with the mutated version with the mutations S:D614G and S:T1117I. The Chimera software (Pettersen et al., 2004) was used not only to generate the mutations in the spike sequence of genomes from the lineage B.1.1.389, but also to minimize energy and visualize the molecules.

Afterward, molecular docking was implemented using the DockThor server (Guedes et al., 2021), in which a grid box parameters were standardized (center x = 221.9245, center y = 208.9935, center z = 195.6175, total size x = 40 Å, total size y = 40 Å, total size z = 40 Å, and discretization = 0.42 Å) in the regions FP, HR1 and HR2 of the spike, similar to (Sixto-López et al., 2021). Comparison between the WT and mutated proteins in the docking analysis was done using energy, in which the conformation with the lowest free energy values was chosen as the most stable for each case.

### Epitope analysis

Spike protein-based epitopes were predicted using IEDB tool (Immune Epitope Database, https://www.iedb.org/), including the following parameters: lineal and discontinuous peptides, for T and B cells as well as MHC ligands, class I and II MHC molecules, human host, and COVID-19 disease. The spike protein sequence (NCBI ID: YP_009724390.1) of the SARS-CoV-2 reference genome (wild type, WT) was used as model. After the prediction was achieved, all the candidate peptides covering the position T1117 in the spike were selected. Then, the modification to I1117 (mutated, S:T1117I) was done manually. Predictions of binding and processing (T and B cells, and the MHC) were re-run with the mutated peptide using specific tools of the Epitope Analysis Resource (part of the IEDB tool). Physicochemical properties and toxicity were evaluated using ToxinPred (Gupta et al., 2013), and allergenicity prediction was done with AllergenFP tool (Dimitrov, Naneva, Doytchinova, & Bangov, 2014). Scores for all the predictions were compared for the WT and mutated peptides.

## Results

### SARS-CoV-2 genomes harboring the mutation S:T1117I define a polyphyletic group with multiple lineages around the world, including the lineage B.1.1.389 as a local genotype

A total of 1155 SARS-CoV-2 sequences were found with the mutation S:T1117I in 54 countries worldwide until April 30^th^, 2020. United States (USA), England, and Costa Rica are the top locations in which this mutation has been reported, with 268 (23.2%), 266 (23.0%), and 126 (10.9%), respectively. However, among all the sequenced genomes for the United States and England, the cumulative prevalence for genomes carrying S:T1117I is <0.5%, whereas for Costa Rica represents 22% of the sequenced cases up to April 30^th^, 2020.

The phylogenetic analysis of these sequences (Figure 1) defines a polyphyletic group, suggesting multiple and independent origins with a divergent profile. This observation is also supported by the diversity of lineages (89 in total), in which 68.7% of genomes belongs to six distinct genotypes: B.1.1.7 (240 sequences, most from European countries), B.1.1.1 (149, most from England), B.1.1.389 (141, most from Costa Rica), B.1 (135, most from USA), B.1.177 (73, most from European countries) and B.1.1 (56, most from England). More details in Supplementary Table S1.

**Figure 1.**
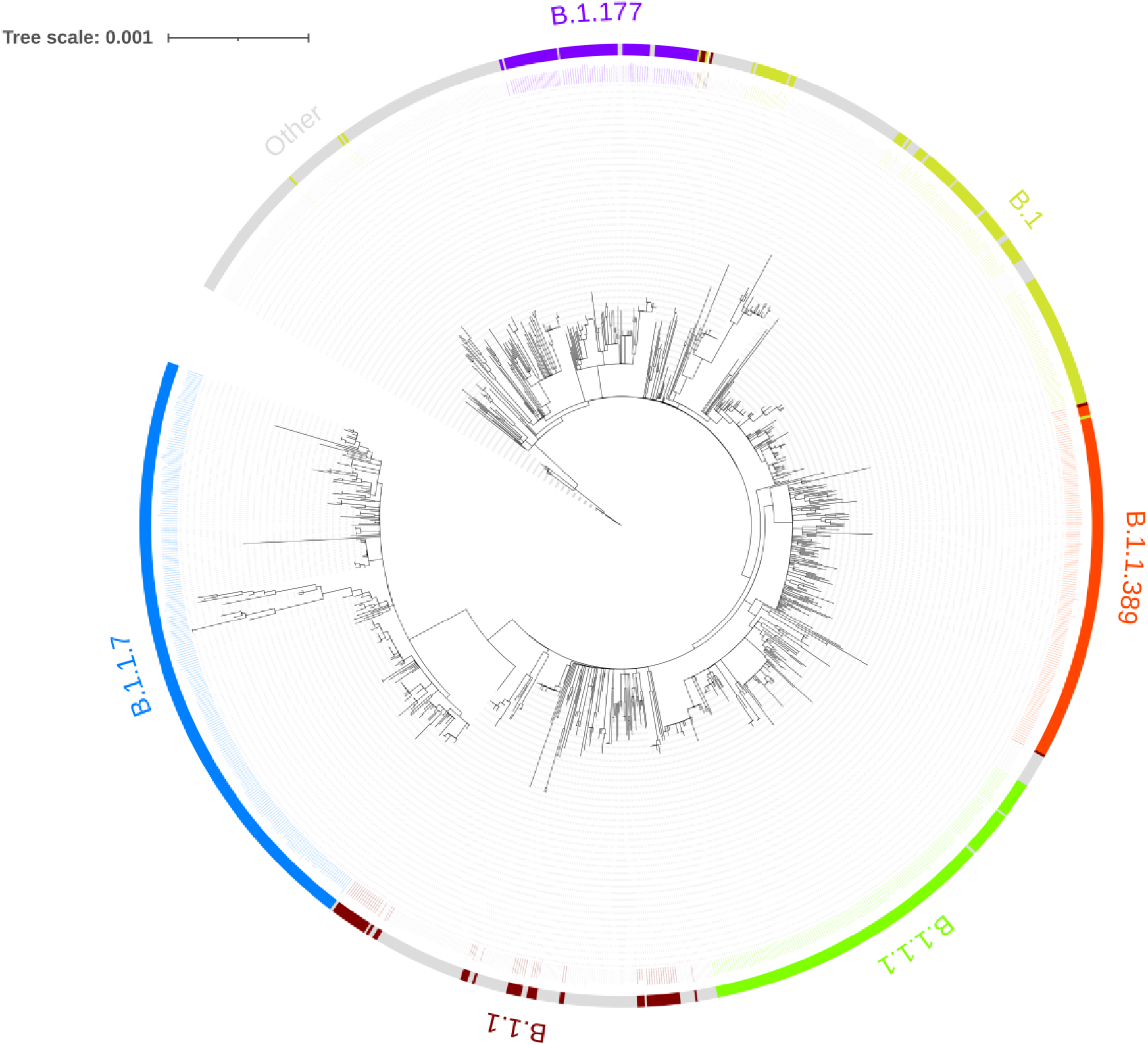
Phylogenetic tree of SARS-CoV-2 genome sequence carrying the mutation T1117 in the spike (S:T1117I) of all around the world. The 1155 available sequences in GISAID database (until April 30^th^, 2021) with this variant are distributed according to PANGOLIN lineages. Six lineages are predominant (frequency >5%), each one with a monophyletic origin.

Due to Costa Rica is part of the top 3 countries reporting more genomes with S:T1117I (with 126 sequences), those genomes are part of the local lineage B.1.1.389, and the last was the predominant group according to the cumulative prevalence in this country during 2020 and early 2021, we followed our analysis studying this genotype and this mutation.

The lineage B.1.1.389 is characterized by the presence of at least eight mutations among the genomes (Figure 2-A), including the D614G and T1117I in the spike. This genotype has been reported in all the seven provinces of Costa Rica with a prevalence range between 10 and 41%, reaching up to 26% out of all sequences (Figure 2-B-C). In addition, new lineages started to circulate during 2021 in Costa Rica, including the A.2.5, A.2.5.1, A.2.5.2, B.1.1.7 (alpha variant) and P.1 (gamma variant), with the subsequent reduction of the B.1.1.389 in the first months of the year 2021 (Figure 2-D).

**Figure 2.**
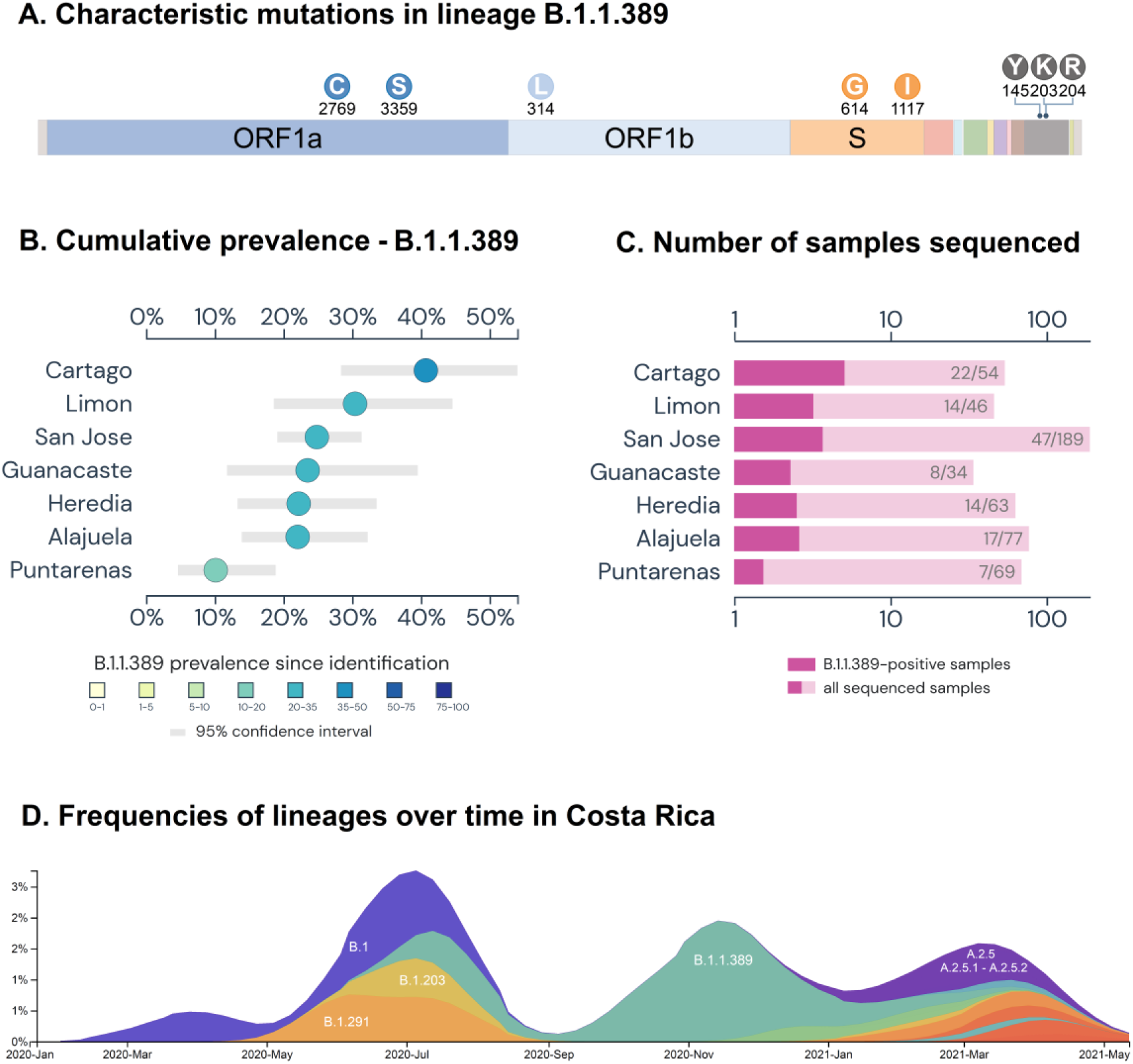
Epidemiological and genomic determinants of the B.1.1.389 lineage. (A) The lineage is characterized by the presence of eight mutations including two in the spike, the D614G and the T1117I variants. (B-C) The B.1.1.389 lineage has been found in all the seven provinces of Costa Rica (prevalence range 10-41%), reaching up to 22% out of all sequences. (D) New lineages started to circulate during 2021 in Costa Rica, including the A.2.5, A.2.5.1, A.2.5.2, B.1.1.7 (alpha variant), and P.1 (gamma variant), with the subsequent reduction of the B.1.1.389.

### Mutation S:T1117I found in the Costa Rican lineage B.1.1.389 is product of natural selection, with some effects on the activity of the function and interactions of the spike protein

In order to analyze the evolutionary context of the mutation S:T1117I and its relation with all the available genomes, we aligned all the 407 spike sequences from SARS-CoV-2 genomes from Costa Rica (with or without the mutation). The reference sequence was also included, completing 408 sequences.

We first studied the natural selection of mutations in the spike protein. The analysis of positive/adaptive selection of mutations among spike sequences found eight sites using the FUBAR model, in which the dN/dS was >1 (Table 1). Tellingly, the sites 501 and 1118, corresponding to the mutations S:N501Y and S:D1118H present in the lineage B.1.1.7, and 1117, site of the mutation of our interest, were included.

**Table 1.**
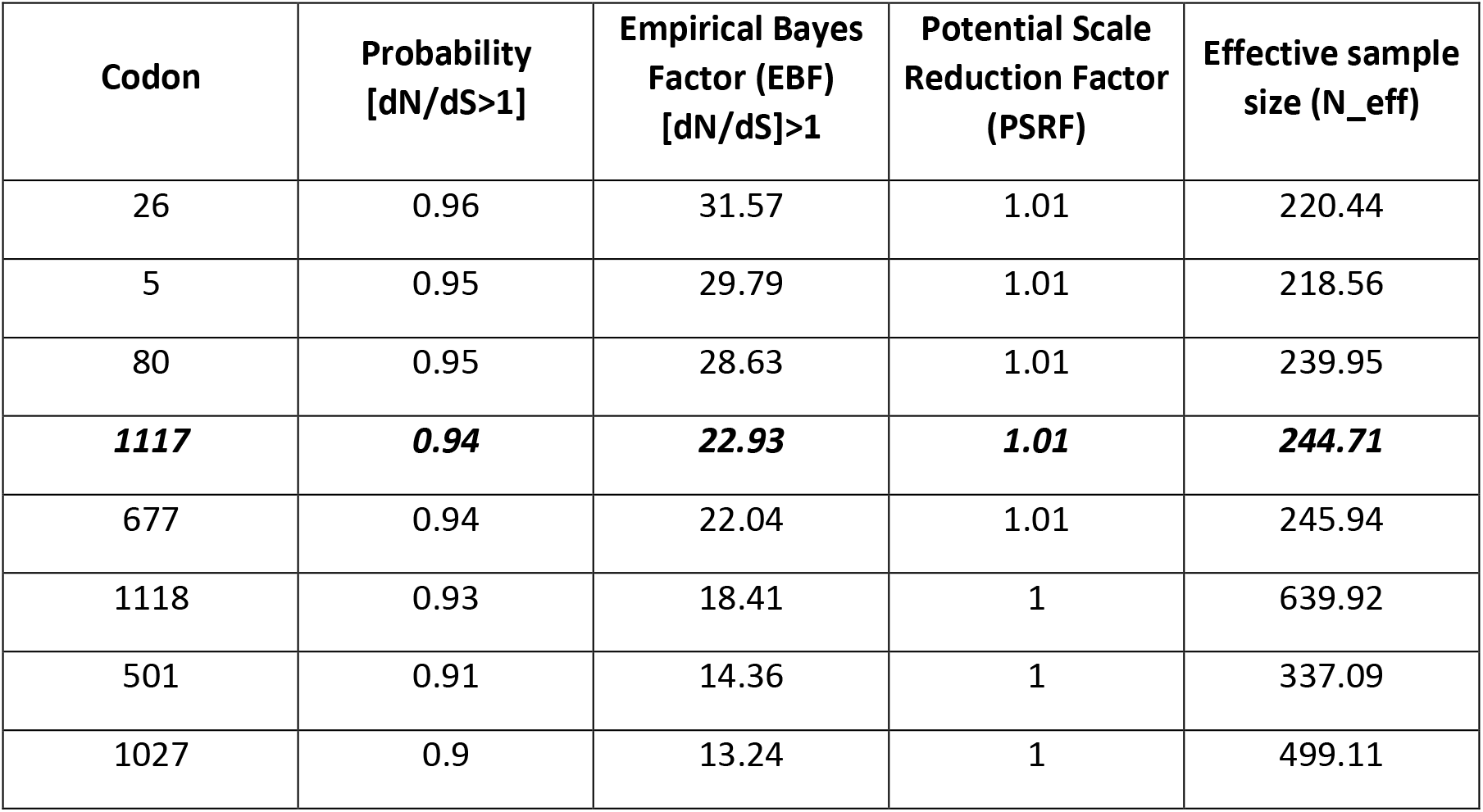
Analysis of positive selection of mutations by a Fast Unconstrained Bayesian AppRoximation (FUBAR) among protein sequences of the spike of the SARS-CoV-2 from Costa Rican cases of COVID-19.

In contrast, as shown in Figure 3, the analysis of residues found no drastic effects of the mutation S:T1117I on the spike in a co-evolutionary context using mutual information and residues conservation models. In this sense, the corrected Mutual Information (MI) was used to identify correlations between positions and the possible effect on the structure or function of the protein, revealing that the most impacted region (orange connections) are part of the RBD (positions 319-541), including the case of the mutation N501Y.

**Figure 3.**
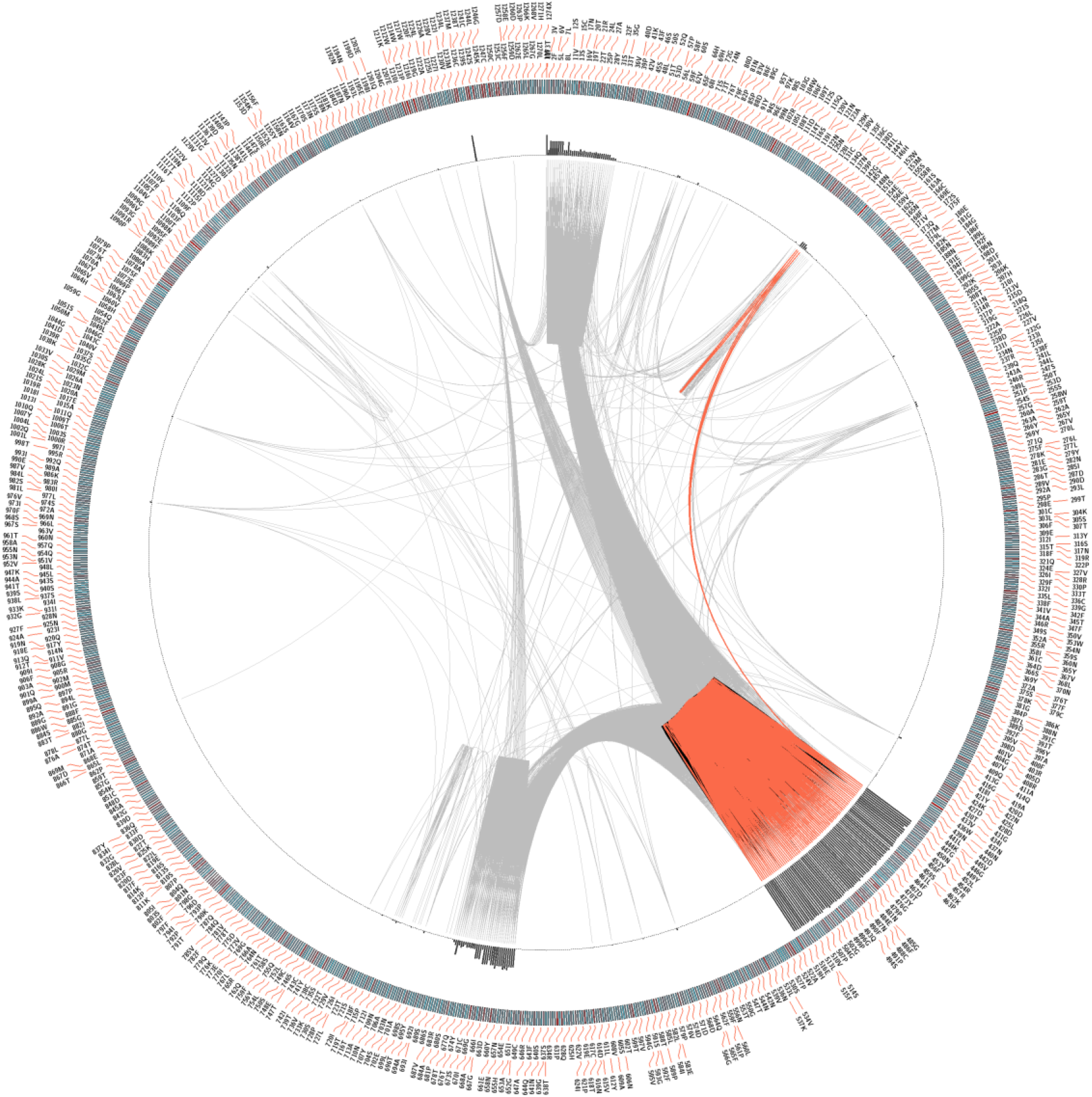
Mutual coevolutionary relationship between residues in the spike protein of SARS-CoV-2 from Costa Rican cases of COVID-19. After a multiple sequence alignment of protein sequences was done, the corrected Mutual Information (MI) was used to identify correlations between positions and the possible effect on the structure or function of the spike protein. The most impacted region (orange connections) are part of the RBD (position 319-541), including the case of the mutation N501Y. For the case of the mutation S:T1117I, no drastic effects are predicted according to the correlation metrics.

Afterward, we continued our study with a molecular docking approach to assess how the mutations in the spike are affecting the molecular interactions. The WT sequence and the mutated version with both mutations S:D614 and S:T1117I (spike sequence for the B.1.1.389 genomes) were considered for the docking with nelfinavir drug as ligand. After modeling, the drug was located in the HR1 region of S2 domain in the spike (Figure 4), as expected. As shown in Table 2, the affinity of the nelfinavir increased because of lower energy was reported for the mutated version (−9.656 kcal/mol) in comparison to the WT spike sequence (−9.231 kcal/mol). The same pattern was observed for total and van der Waals energies, unlike the electrostatic energy.

**Table 2.**
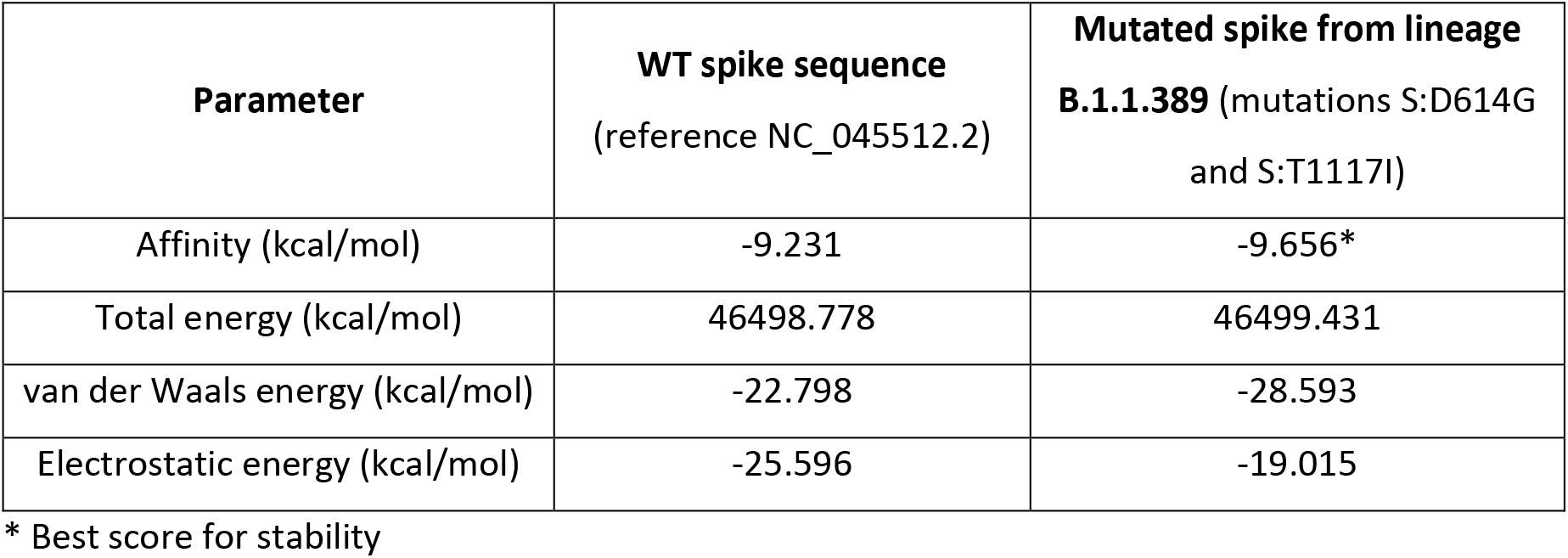
Molecular docking between nelfinavir and the spike protein of the SARS-CoV-2 for WT and the mutated sequences.

**Figure 4.**
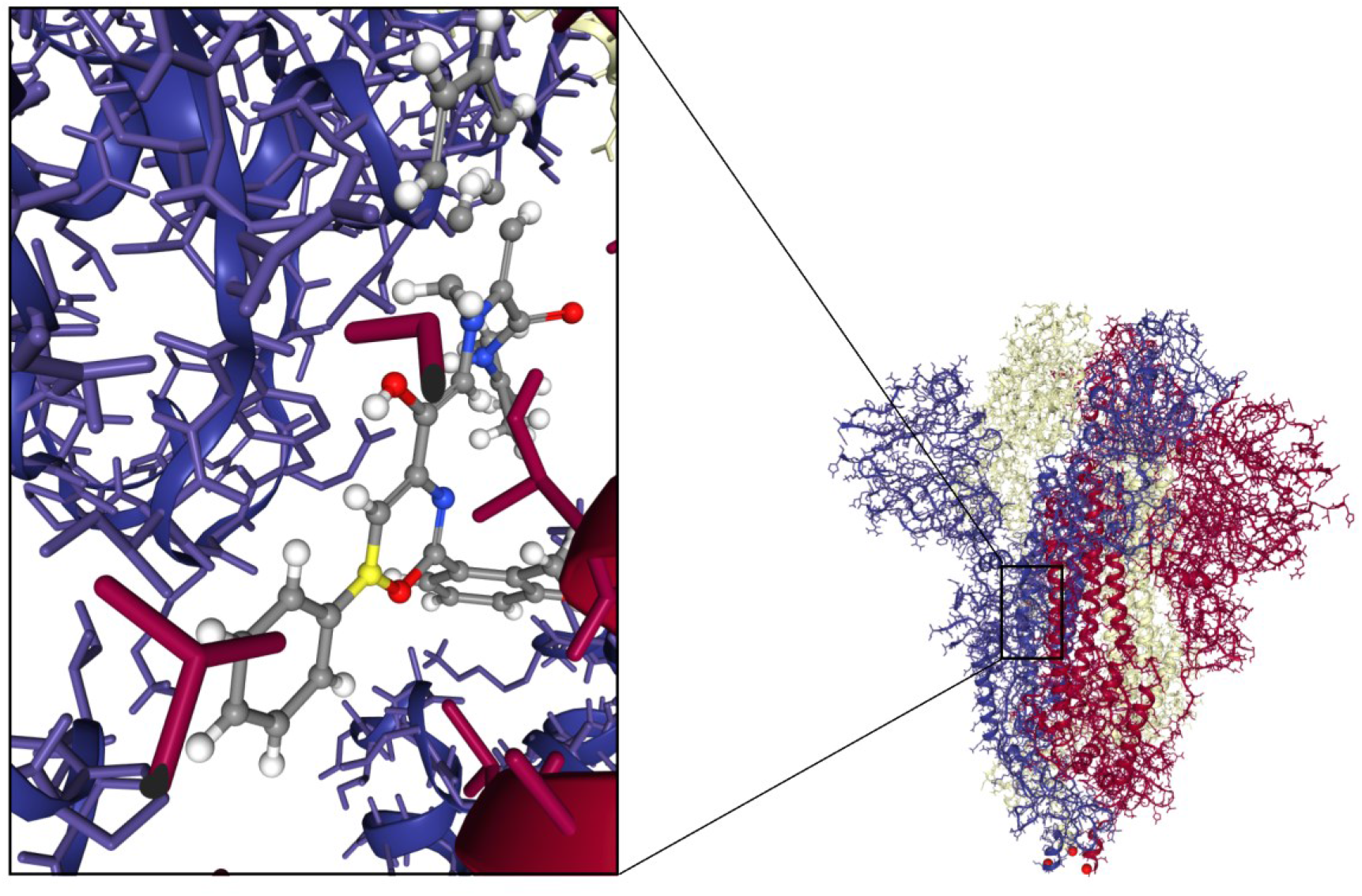
Molecular docking of the complex protein-ligand using the mutated spike protein (lineage B.1.1.389) and nelfinavir drug. The drug was docked into the HR1 region of S2 domain in the spike, using the WT or the mutated (S:D614G and S:T1117I) proteins. Affinity energy predicted a more stable complex for the mutated one.

Finally, to describe the effect of the mutation S:T1117I on the immune activity, an immunoinformatic approach to study peptides was performed. Epitopes were predicted for the spike protein, including lineal or continuous peptides able to bind and be processed by T and B cells, and MHC molecules. Results are shown in Table 3. A total of 742 candidate peptides were identified but only one epitope overlapped the 1117 position (IEDB-ID: 1309561), which corresponded to a linear B cell peptide. The comparison of the WT and the mutated version of the sequence showed changes in physicochemical properties, but scores of recognition by B cells and binding to MHC molecules resulted more favorable for the WT peptide, unlike the processing by MHC-I.

**Table 3.**
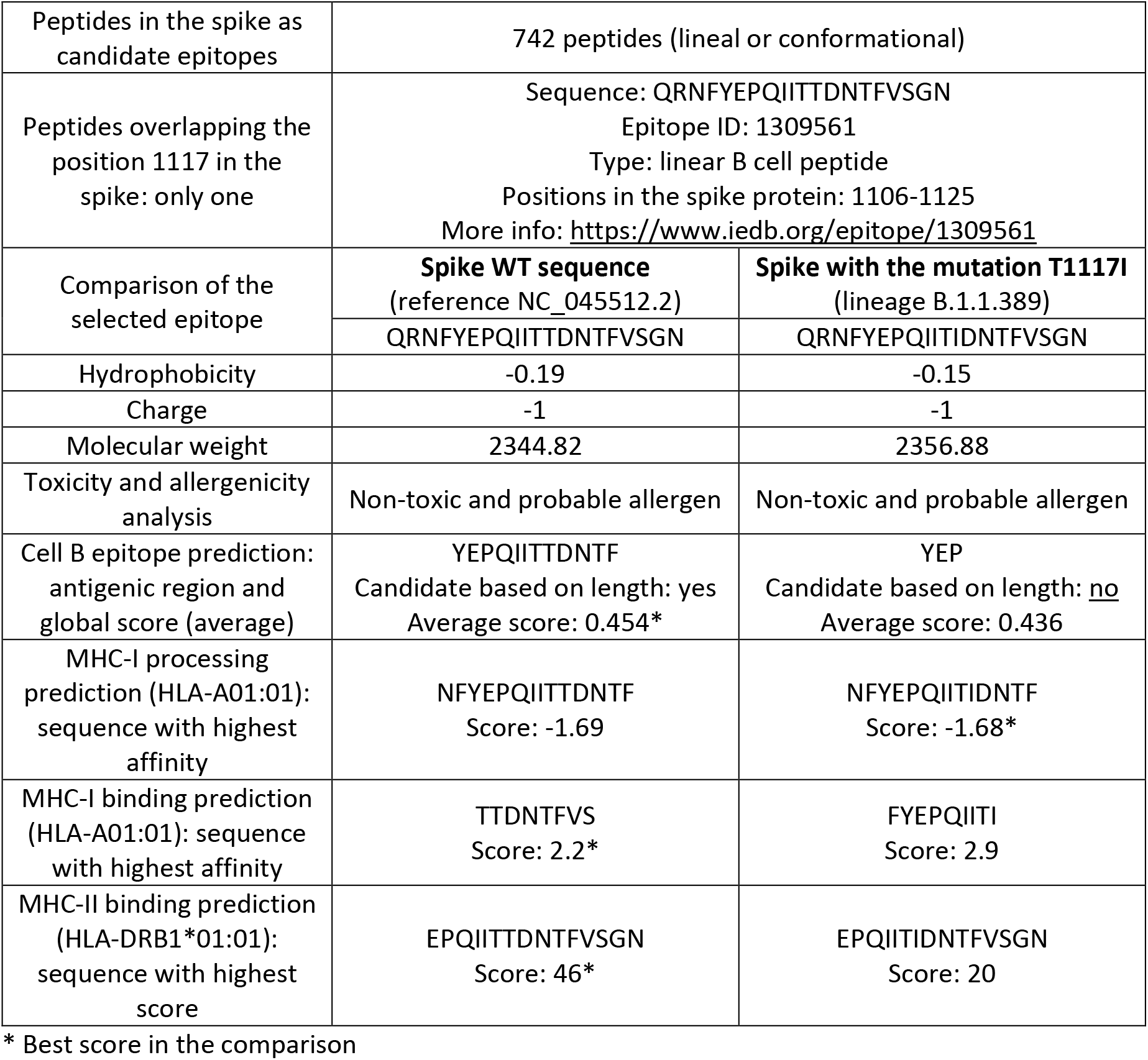
Comparison of epitopes associated with the mutation S:T1117I of the SARS-CoV-2.

## DISCUSSION

Genome sequencing has been a primordial step in the surveillance of the SARS-CoV-2 virus to identify mutations and new genotypes (Castillo, Parra, Tapia, Acevedo, et al., 2020; Graham et al., 2021). According to GISAID database, more than 2 million cases have been sequenced around the world until June 30^th^, 2021.

With the emergency of new variants, the characterization of mutations is relevant because the virus can change transmission, severity, mortality, vaccine effectiveness, and others (Graham et al., 2021; Mercatelli & Giorgi, 2020; Tegally et al., 2020). In Costa Rica, the alpha (lineage B.1.1.7), beta (B.1.351) and gamma (P.1) VOCs have been reported in the first semester of the year 2021 (https://www.gisaid.org/), unlike the delta (B.1.617.2). However, during 2020, lineage B.1.1.389 was predominant in this country, with an accumulative prevalence of 30% by January 2021 (Molina-Mora, Cordero-Laurent, et al., 2021), and up to 22% by April 2021 (this study). Owed to the key mutation of the B.1.1.389 is the S:T1117I, we thereby analyzed to study the effects of this change on the spike function.

We started by comparing all the available sequences with the S:T1117I around the world. The 1155 sequences defined a polyphyletic tree using phylogeny, in which 89 distinct lineages were recognized, including six dominant ones. Most of the cases were identified in the USA, England, and Costa Rica, although only the last had a high cumulative prevalence of 22%, in contrast to the <0.5% for the others. Similar patterns with polyphyletic groups and different prevalence have been also reported for other mutations, including studies in other Latin American countries (Candido et al., 2021; Castillo, Parra, Tapia, Lagos, et al., 2020; Ferrareze et al., 2021; Muñoz, Patiño, Ballesteros, Paniz-Mondolfi, & Ramírez, 2021; Taboada et al., 2020).

Afterward, we followed the analysis using natural selection models, with all the available spike sequences from Costa Rica, to determine the evolutionary role of mutations in the spike protein. Eight sites or mutations were under positive/adaptive selection, including S:T1117I. Interestingly, S:N501Y and S:D1118H from lineage B.1.1.7 (alpha variant) were also recognized. In general, these results are in line with other similar approaches worldwide, in which have been demonstrated that spike protein is under persistent positive selection (Berrio, Gartner, & Wray, 2020; Ferrareze et al., 2021; Lythgoe et al., 2021). Our analysis is the first one with the report of selection for the S:T1117I. Nevertheless, a different scenario for the S:T1117 was found using mutual information to study positions and the possible effect on structure or function. The investigation of the co-evolutionary context of the spike mutations found that the RBD is the most impacted region by the mutations, as may be expected. No drastic effects were recognized for the region associated with the mutation S:T1117I. Only a few works have implemented this approach to study mutations of SARS-CoV-2, in which a reduction of the conservation for the RBD region was also demonstrated, with implications in the interaction of the spike with antibodies (G. Verkhivker, 2020; G. M. Verkhivker & Di Paola, 2021). Even though the natural selection models and the co-evolutionary approach are distinct approaches, the former is based on stochastic evolutionary models only using sequence data (Kosakovsky Pond et al., 2005), while the second one uses sequences and the structural model of the spike (Colell et al., 2018; Simonetti et al., 2013), which could explain the observed results.

On the other hand, according to the architecture of the spike protein (Xia et al., 2020), the S:T1117I is located between the HR1 and HR2 regions of the S2 domain. Preliminary predictions of the effects of this mutation on the function of the spike were suggested in the oligomerization needed for cell infection (Molina-Mora, Cordero-Laurent, et al., 2021). To gain more insights into the relevance of the mutation, molecular docking was performed using nelfinavir drug against the spike protein. For both, the WT and the mutated (S:D614 and S:T1117I) spike proteins, the drug was docked in the HR1 region of S2 domain, as reported before (Musarrat et al., 2020). Although the mutations are far from the RBD region, it is known that outside binding site mutations affect the ligand recognition (Sixto-López et al., 2021), as found here with changes in affinity energy. A higher affinity was found for the mutated spike in comparison to the WT version, with energy values that are in line with other studies including WT and mutations in the spike protein using nelfinavir (Musarrat et al., 2020; Sixto-López et al., 2021) or other drugs (Calligari, Bobone, Ricci, & Bocedi, 2020; Hall & Ji, 2020). These results suggest that the mutated spike sequence defines a more stable complex with the ligand than the WT protein and it could be more affected by the inhibition mechanism of the drug, as previously reported for other mutations in the spike for this drug and other molecules (Sixto-López et al., 2021; G. M. Verkhivker & Di Paola, 2021).

Finally, we analyzed antigenic peptides from spike protein that could be affected by the mutation S:T1117I using immunoinformatics, similar to other approaches (Li et al., 2020; Lin et al., 2020). A total of 742 epitopes were predicted for the spike protein, but only one overlap the site of the mutation. The analysis of the effect of the mutation for that sequence revealed the best scores for the WT sequence, with some changes in the physicochemical properties. These results shows that the S:T1117I changes the properties of the epitope to be recognized by the immune system. However, the peptide is located out of the RBD (S1 domain) and, more importantly, the peptide corresponds to only one peptide out of 742 candidates that are predicted to induce an immune response, suggesting that the vaccination effectiveness is not affected by this mutation.

Altogether, the different effects of the mutation were found on the function of the spike protein, the interaction with molecules, and immunity, showing a complex scenario regarding the real value of this mutation in the spread of SARS-CoV-2 and the pandemic. Such a scenario and the fact that the lineage B.1.1.389 was predominant during the second semester of 2020 could suggest that the mutation represented a partial advantage in that period. Nevertheless, this benefit could be no strong enough with the introductions of other lineages such as the lineages A.2.5, A.2.5.1 and, A.2.5.2 (with high frequency in Central America) which were predominant during 2021, and later with the arrival of the alpha and gamma variants. In addition, our previous work demonstrated that the clinical manifestations of COVID-19 in Costa Rican cases were independent of the SARS-CoV-2 lineages, including no differences for the lineage B.1.1.389 and the mutation S:T1117I (Molina-Mora, González, et al., 2021).

COVID-19 disease, as an infection disease, depends not only on the viral agent (SARS-CoV-2 genotype), but also on the host conditions (genetics and risk factors) and the environment (social behavior) (Molina-Mora, González, et al., 2021; Tsui, Deng, & Pan, 2020). Thus, this work represents another piece to understand the dynamics of the COVID-19 pandemic in Costa Rica, in this case focused on the virus. Further analyses are required to validate experimentally predictions and results, which is the main limitation of this *in silico* study. Furthermore, this work offers a pipeline using distinct bioinformatics analyses, which can be applied to other mutations in the spike, proteins of SARS-CoV-2, or pathogens.

## Conclusions

In conclusion, this work analyzed the mutation T1117I in the spike protein of the SARS-CoV-2. Insights into the biological meaning of this change were gained, including the description of a polyphyletic pattern around the world, with the emergency of local lineages. In Costa Rica, the mutation is found in the lineage B.1.1.389 and it is suggested it was generated as a product of positive/adaptive selection. Different changes in the function of the spike protein, higher affinity in the interaction with molecules, and scarce changes in immunity were revealed for this mutation. This suggests a partial benefit for the spread of the virus with this genotype during the year 2020 in Costa Rica, although possibly not strong enough with the introduction of new lineages during early 2021 which became predominant later. In addition, the bioinformatic strategy developed here can be eventually used to analyze other mutations using *in silico* approaches.

## Ethical approval and consent to participate

This study was approved by the scientific committee of CIET-UCR (No. 242-2020). Consent was not required for this study.

## Acknowledgments

We thank all the researched who sheared all sequences and other data into public databases, including the public laboratories from Costa Rica, as well as Meriyeins Sibaja-Amador and Carlos Martínez-Calderón for their assistance in different activities of the project.

## Availability of data and material

Data were retrieved from public databases, as described in Methods. See Supplementary Material for details of sequence ID, locations, lineages and other data.

## Author contributions

The conception and design of the study, data acquisition, bioinformatics analysis, interpretation of results, and writing of the manuscript was done by JMM.

## Funding

This work was funded by Vicerrectoría de Investigación – Universidad de Costa Rica, with the Project “C0196 Protocolo bioinformático y de inteligencia artificial para el apoyo de la vigilancia epidemiológica basada en laboratorio del virus SARS-CoV-2 mediante la identificación de patrones genómicos y clínico-demográficos en Costa Rica (2020-2022)”.

---

**Table S1.**
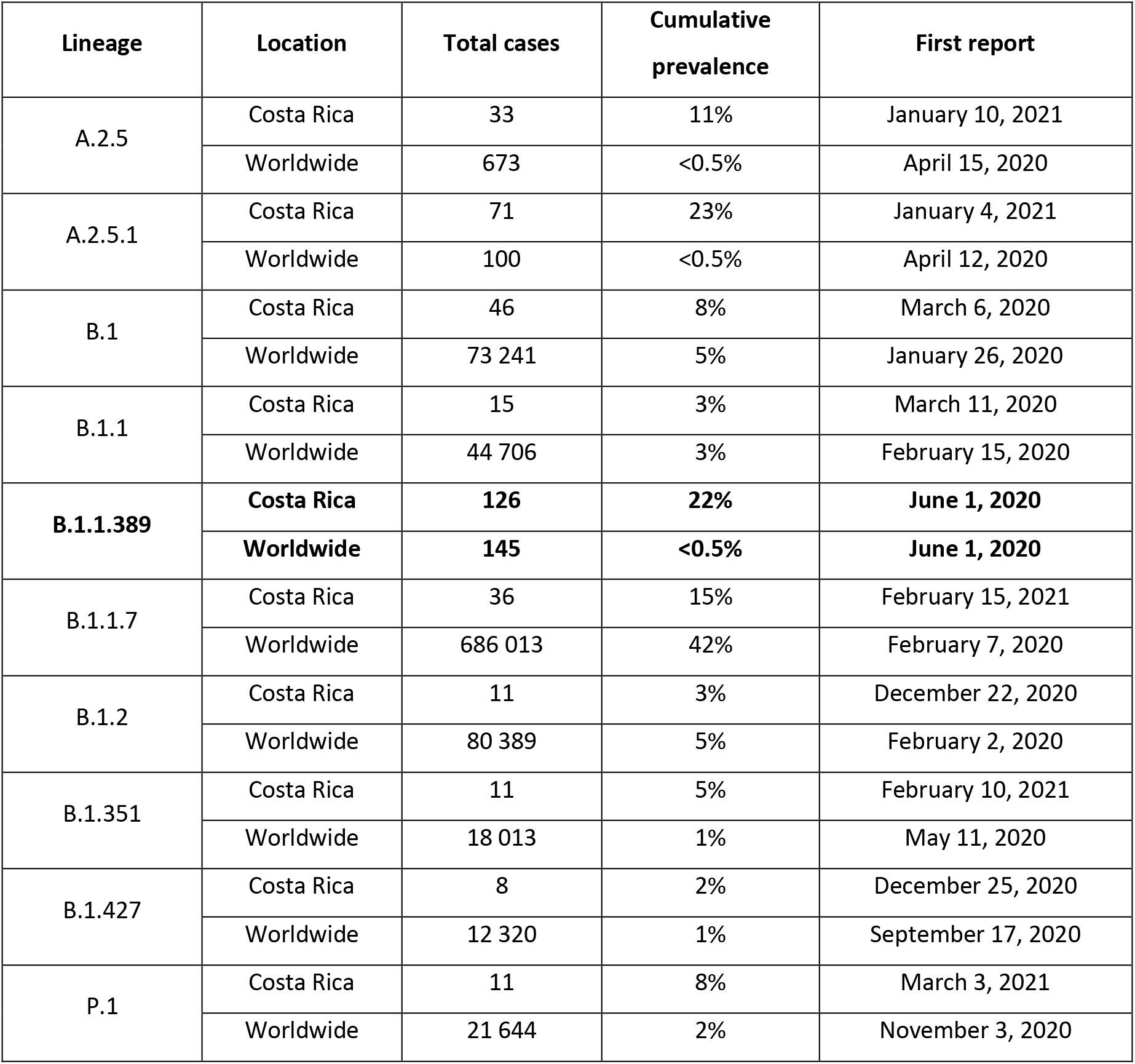
Distribution of SARS-CoV-2 genomes by lineage circulating in Costa Rica until April 30^th^, 2021.

