## Supplementary Table S1 for "Insights into the mutation T1117I in the spike and the lineage B.1.1.389 of SARS-CoV-2 circulating in Costa Rica"

**Table S1. Distribution of SARS-CoV-2 genomes by lineage circulating in Costa Rica until April 30^th^, 2021.**

| **Lineage** | **Location** | **Total cases** | **Cumulative prevalence** | **First report** |
| --- | --- | --- | --- | --- |
| A.2.5 | Costa Rica | 33 | 11% | January 10, 2021 |
|  | Worldwide | 673 | <0.5% | April 15, 2020 |
| A.2.5.1 | Costa Rica | 71 | 23% | January 4, 2021 |
|  | Worldwide | 100 | <0.5% | April 12, 2020 |
| B.1 | Costa Rica | 46 | 8% | March 6, 2020 |
|  | Worldwide | 73 241 | 5% | January 26, 2020 |
| B.1.1 | Costa Rica | 15 | 3% | March 11, 2020 |
|  | Worldwide | 44 706 | 3% | February 15, 2020 |
| **B.1.1.389** | **Costa Rica** | **126** | **22%** | **June 1, 2020** |
|  | **Worldwide** | **145** | **<0.5%** | **June 1, 2020** |
| B.1.1.7 | Costa Rica | 36 | 15% | February 15, 2021 |
|  | Worldwide | 686 013 | 42% | February 7, 2020 |
| B.1.2 | Costa Rica | 11 | 3% | December 22, 2020 |
|  | Worldwide | 80 389 | 5% | February 2, 2020 |
| B.1.351 | Costa Rica | 11 | 5% | February 10, 2021 |
|  | Worldwide | 18 013 | 1% | May 11, 2020 |
| B.1.427 | Costa Rica | 8 | 2% | December 25, 2020 |
|  | Worldwide | 12 320 | 1% | September 17, 2020 |
| P.1 | Costa Rica | 11 | 8% | March 3, 2021 |
|  | Worldwide | 21 644 | 2% | November 3, 2020 |
